# Investigation of *Trypanosoma-*induced vascular damage sheds insights into *Trypanosoma vivax* sequestration

**DOI:** 10.1101/2023.09.18.558284

**Authors:** Sara Silva Pereira, Daniela Brás, Teresa Porqueddu, Ana M. Nascimento, Mariana De Niz

## Abstract

Multiple blood-borne pathogens infecting mammals establish close interactions with the host vascular endothelium as part of their life cycles. In this work, we investigate differences in the interactions of three *Trypanosoma* species: *T. brucei, T. congolense* and *T. vivax* with the blood vasculature. Infection with these species results in vastly different pathologies, including different effects on vascular homeostasis, such as changes in vascular permeability and microhemorrhages. While all three species are extracellular parasites, *T. congolense* is strictly intravascular, while *T. brucei* is capable of surviving both extra- and intravascularly. Our knowledge regarding *T. vivax* tropism and its capacity of migration across the vascular endothelium is unknown. In this work, we show for the first time that *T. vivax* parasites sequester to the vascular endothelium of most organs, and that, like *T. congolense, T. vivax* Y486 is largely incapable of extravasation. Infection with this parasite species results in a unique effect on vascular endothelium receptors including general downregulation of ICAM1 and ESAM, and upregulation of VCAM1, CD36 and E-selectin. Our findings on the differences between the two sequestering species (*T. congolense* and *T. vivax*) and the non-sequestering, but extravasating, *T. brucei* raise important questions on the relevance of sequestration to the parasite’s survival in the mammalian host, and the evolutionary relevance of both sequestration and extravasation.

## Introduction

Blood vessel endothelium forms a semi-permeable barrier between the blood contents and the extracellular matrix of tissues. Many blood-borne pathogens, including a plethora of parasites, establish close interactions with the vascular endothelium as part of their life cycles in the mammalian hosts. These interactions are important not only for parasite survival (e.g. allowing immune evasion and/or clearance from the bloodstream), but also key for our understanding of parasite tropism and host pathology. Parasite-host vascular interactions have been a topic of great interest, particularly with respect to protozoan parasites (Bentivoglio et al., 2018; Bernabeu et al., 2021; Bush et al., 2021; De Niz et al., 2021, 2016; Formaglio et al., 2023; Hopp et al., 2015; Mabille et al., 2022; Peters et al., 2008; Ross et al., 2021; Silva Pereira et al., 2022, 2019). The interaction of parasites with the host vascular endothelium is generally linked with major changes in both the host and the parasite, such as altered expression of specific receptors that facilitate pathogen binding to the vascular endothelium; the increase in vascular permeability, which facilitates cell trafficking across the endothelial barrier; or the generation of microhemorrhages that result in organ dysfunction (reviewed in Konradt and Hunter, 2018).

African trypanosomes (*Trypanosoma brucei*, *T. congolense*, *T. vivax*) cause animal African trypanosomiasis in livestock and are a considerable source of animal morbidity and mortality, as well as socio-economic impairment in the tropics. Trypanosome infections can range from asymptomatic to acute, although the most common presentation in cattle is a chronic, wasting disease. The behavior of each parasite species in mammalian hosts is poorly studied, but evidence accumulates supporting that it may directly impact pathology and disease severity. *T. brucei* is known to proliferate in both blood and the extravascular spaces of tissues, which it reaches by crossing the vascular endothelium (Caljon et al., 2016; Capewell et al., 2016; De Niz et al., 2021; Mabille et al., 2022; Masocha et al., 2007; Namayanja et al., 2017; Trindade et al., 2016) by mechanisms not yet fully understood. In contrast, *T. congolense* cytoadheres to the vascular endothelium likely via host sialic acid binding (Hemphill et al., 1994; Hemphill and Ross, 1995), but does not invade tissues (reviewed in Silva Pereira et al., 2019). This mechanism, known as sequestration, was recently shown to determine disease severity in a mouse model of acute trypanosomiasis (Silva Pereira et al., 2022).

Despite these recent advances, the behavior of *T. vivax* parasites in the mammalian host remains elusive. Previous work has described the pathology of *T. vivax* in mouse and livestock models, as well as in natural infections. Findings in goats include evaluation of clinal and pathological manifestations in the central nervous system of infected goats, some of which developed meningoencephalitis and meningitis (Batista et al., 2011). descriptions of parasite presence in the skin and adipose tissue, together with mononuclear inflammatory reactions (Machado et al., 2021), and follicular degeneration in ovaries, together with enlarged lymph nodes and weight loss (Rodrigues et al., 2013). Findings in sheep include descriptions of acute testicular degeneration and hyperplasia of the epididymal epithelium (Bezerra et al., 2008). Evaluation of multi-organ histopathology in mice, includes findings of multifocal inflammatory infiltrates across organs, extramedullary hematopoiesis in the liver, and cerebral oedema (Chamond et al., 2010), as well as observation of parasite infiltration to the brain, liver and lungs (D’Archivio et al., 2013). Investigations in natural infections in calves and goats showed systemic symptoms including depression, weight loss, pale mucous membranes, enlarged lymph nodes, oedema, cough, coryza and diarrhoea (Batista et al., 2012, 2009), as well as blindness and abortion (Batista et al., 2009); and gross lesions to the meninges, lymph nodes and spleen (Batista et al., 2007).

Altogether, however, whether *T. vivax* can display tropism to and establish in tissues has been long debated (Machado et al., 2021 and reviewed in Silva Pereira et al., 2019). Here, we use intravital and *ex vivo* fluorescence microscopy to compare the organ distribution of each of the three aforementioned trypanosome species in mouse models, the vascular changes induced by them, and their impact on the expression of key blood vascular endothelial cell receptors. We show for the first time that *T. vivax* (strain Y486) cytoadheres to the vascular endothelium, albeit differently from *T. congolense*, but we found no evidence of tissue extravasation during the early stage of infection up to the first peak of parasitemia. This work highlights the impact of trypanosome-vascular interactions for the host and suggests that sequestration might be a trait carried over from the last common ancestor of African trypanosomes.

## Results

### *T. brucei, T. congolense* and *T. vivax* infections result in different clinical presentations

It has long been known that despite common nomenclature, different African trypanosome species cause different diseases, with widely disparate symptomatology. However, the specificities of the host response to each *Trypanosoma* species are still poorly understood. Here, we began by comparing how infection with a well characterized *T. brucei* AnTat fluorescent reporter line (Calvo-Alvarez et al., 2018), *T. congolense* savannah 1/148 (MBOI/NG/60/1–148) (Young and Godfrey, 1983), and *T. vivax* Y486 (Gibson, 2012) progressed in C57BL/6 mice. The infection progression in C57BL/6 mice with these *T. brucei* and *T. congolense* lines has been previously described (De Niz et al., 2021; Silva Pereira et al., 2022). Briefly, infection with a medium (10^4^ parasites/ml) inoculum of *T. brucei* results in approximately a 20-day-long infection (De Niz et al., 2021) comprising two to four peaks of parasitaemia (**Figure 1A-1B, blue lines**). *T. congolense* infection results in acute symptomatology that is lethal by the first peak of parasitaemia around day 7 post-infection (p.i.) (Silva Pereira et al., 2022) (**Figure 1A-1B, green lines**). In contrast, *T. vivax* Y486 results in either a short infection (mice dying between before day 11 p. i.) or a chronic infection that can extend to 60 days, characterized by several peaks of parasitaemia (**Figure 1A-1B, red lines**). The median survival is 20 days for *T. brucei*, 8.5 days for *T. congolense*, and 28 days for *T. vivax* (p-value < 0.0001, Gehan-Breslow-Wilcoxon test). Additional to the different parasitemia profiles, symptoms differed widely amongst the 3 species, with *T. brucei*-infected mice showing significant cachexia and difficulty breathing at the time of death. Conversely, *T. congolense*-infected mice showed symptoms consisting of neurological damage, liver damage and cachexia. Finally, close to the time of death, *T. vivax*-infected mice show signs of overt disease, including apathy, difficulty breathing, weakness, but no obvious weight loss or cachexia.

**Figure 1.**
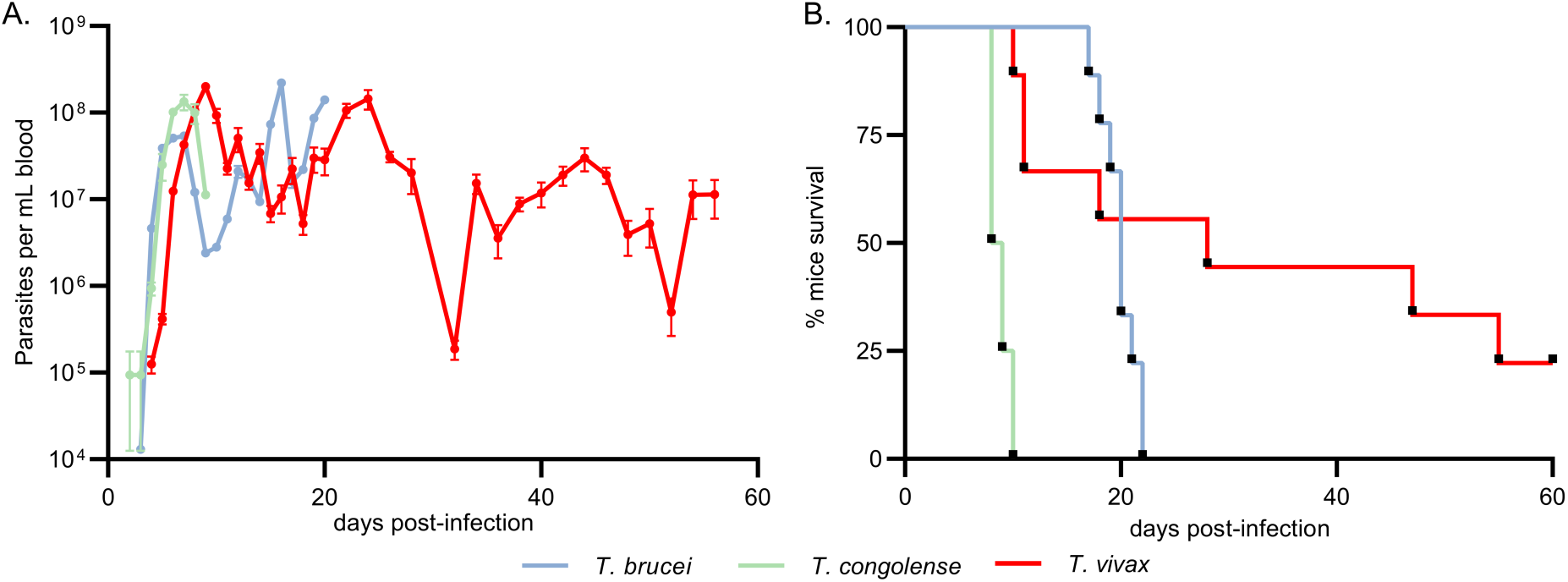
Infection progression of African trypanosomiasis in C57B/6J mice. A. Parasitemia throughout infection with *T. brucei* Antat 1.1, *T. congolense* 1/148, and *T. vivax* Y486 parasites, estimated by hemocytometry. Mean ± SEM. B. Mice survival curves following infection with *T. brucei* Antat 1.1, *T. congolense* 1/148, or *T. vivax* Y486 (median survival of 20, 8.5, 28 days post-infection, respectively, p-value < 0.0001, Gehan-Breslow-Wilcoxon test comparing survival curves) (N=4-10). Data from *T. brucei* and *T. congolense* infections was reused from De Niz *et al*. 2021 and Silva Pereira *et al*. 2022, respectively.

### Infection-induced vascular changes are species- and tissue-dependent

Following the characterization of parasitemia progression, we analyzed the effect of infection with each *Trypanosome* species on vascular homeostasis, measured by vascular permeability and the presence of hemorrhagic lesions until the first peak of parasitaemia (defined as day 7 p.i. for *T. brucei,* day 6 for *T. congolense,* and day 8 for *T. vivax*). We measured vascular permeability using previously described methodology (De Niz et al 2021, Silva-Pereira et al 2022), namely, intravenously injecting FITC-Dextran 70kDa followed by time-lapse intravital microscopy to detect fluorescence intensity changes in the intra- and extra-vascular compartments in each organ, at each day post-infection (**Figure 2A**). Upon comparison of the 3 species, we detected a widely different vascular pathology profile in terms of time of the initiation of pathology, organs affected, and maximum vascular pathology detected. During *T. brucei* infection, the spleen was the first organ to display increased permeability at day 3 p. i., followed by the adipose tissue from day 4 p. i., and the pancreas, liver and kidney, from day 5 p. i. (**Figure 2B blue lines, 2C top panel**). During the first peak of parasitemia, a maximum increase in vascular permeability was detected in the adipose tissue (with a 6.7-fold increase relative to control mice) followed by the liver, spleen and pancreas, reaching a c.a. 5-fold increase in permeability relative to control. These results were previously discussed in detail in (De Niz et al., 2021). Upon *T. congolense* infections, vascular permeability increased earlier than in *T. brucei* infections, more organs were affected than for *T. brucei* during the first parasitemia peak, and a higher fold-change in permeability was recorded for all organs compared to *T. brucei* infections. Namely, from day 1 p. i., vascular permeability increased in the heart; from day 2 p. i. in the liver, from day 3 p. i. in the lungs, spleen, and pancreas; from day 4 p. i. in the adipose tissue, and from day 5 p. i. in the kidney (**Figure 2B green lines, 2C middle panel**). It is worth noting that vascular permeability in the liver, heart and spleen, increased by between 10- and 15-fold by day 6 p. i. Finally, *T. vivax* infection resulted in milder and later changes in permeability than either of the other two species. Overall, the heart showed increased permeability from day 2 p. i., the adipose tissue from day 5 p. i., and the lungs right at the peak of parasitaemia (day 8 p. i.). The maximum fold-change in vascular permeability was detected in the adipose tissue (9-fold increase relative to control) followed by the heart and lungs (between 6- and 8-fold increase relative to control) (**Figure 2B red lines, 2C bottom panel**). Consistent with the physiology and tight regulation of the blood brain barrier, the brain was the organ with the lowest change in vascular permeability throughout infection with the three species (**Figure 2B, top left panel**), despite the presence of hemorrhagic lesions, particularly during *T. congolense* infection.

**Figure 2.**
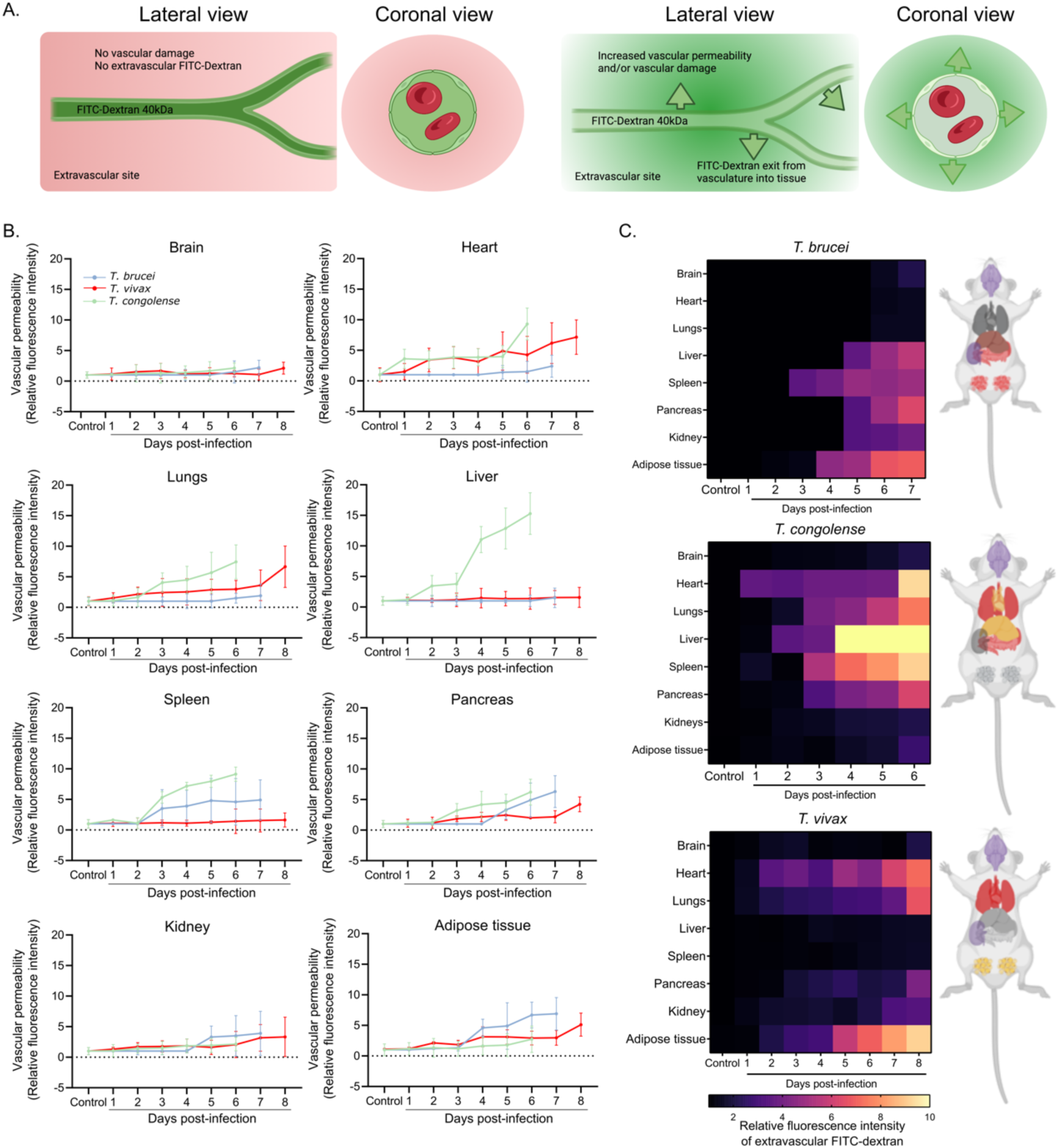
Dynamics of vascular permeability during African trypanosomiasis in C57B/6J mice. A. Schematic representation of vascular permeability assessment. FITC-Dextran 70kDa was administered intravenously to mice. If there is no vascular damage and permeability is not compromised, FITC-Dextran remains intravascular. If there is vascular damage and/or increased vascular permeability, FITC-Dextran diffuses to the extravascular spaces and can be detected by fluorescence microscopy. B. Vascular permeability of major organs upon infection with *T. brucei*, *T. congolense*, or *T. vivax*, during the first peak of parasitaemia. Mean fluorescence intensity of extravascular FITC-Dextran relative to non-infected mice is used as a proxy for vascular permeability. C. Heatmaps and respective anatomical diagrams showing data presented in B, color-coded according to key. Data from *T. brucei* and *T. congolense* (brain only) infections was reused from De Niz *et al*. 2021 and Silva Pereira *et al*. 2022, respectively.

We define vascular permeability changes and hemorrhagic lesions as two different phenomena: with the former referring to increased FITC-Dextran leakage to the organ’s extravascular space, and the latter being extravascular presence of red blood cell aggregations resulting from a physical disruption of the vessel wall. We identified micro-hemorrhages by the accumulation of FITC-Dextran and the presence of red blood cells in extravascular regions (**Figure 2A** and **Figure 3**). We defined 4 categories of damage as follows: category 1: no hemorrhages and no increased permeability (grey); category 2: no hemorrhages and increased permeability (pink); category 3: hemorrhages and no increased permeability (light red); category 4: hemorrhages and increased permeability (dark red) (**Figure 3**). Images shown in **Figure 3** correspond to the peak of parasitemia with each species. Results for *T. brucei* were previously published in (De Niz et al., 2021). The number of vessels with hemorrhagic lesions was considerably higher in *T. congolense* infections than the remaining species, at the first peak of parasitaemia, consistent with previously described acute cerebral disease (**Figure 3**) (Silva Pereira et al., 2022). Hemorrhagic lesions were present in most organs, regardless of the trypanosome species, albeit at different incidence and severity levels. *T. congolense* infection resulted in the highest level of hemorrhages throughout major organs (liver (88%), spleen (47%), brain (33%), pancreas (32%), lungs (30%), adipose tissue (23%), heart (22%), and kidney (20%)). Conversely, *T. brucei* and *T. vivax* infections resulted in overall lower levels of hemorrhages, although still detectable in major organs (in *T. brucei*: kidney (29%), spleen (17%), brain (15%), liver (12%), pancreas (8%), lungs (7%), and adipose tissue (4%); in *T. vivax*: (28%), spleen (15%), kidney (11%), brain (10%), lungs (10%), adipose tissue (10%), heart (8%), and pancreas (4%)). These results suggest that African trypanosomes induce species- and tissue-specific vascular changes that range from increased vascular permeability to vascular micro-hemorrhages.

**Figure 3.**
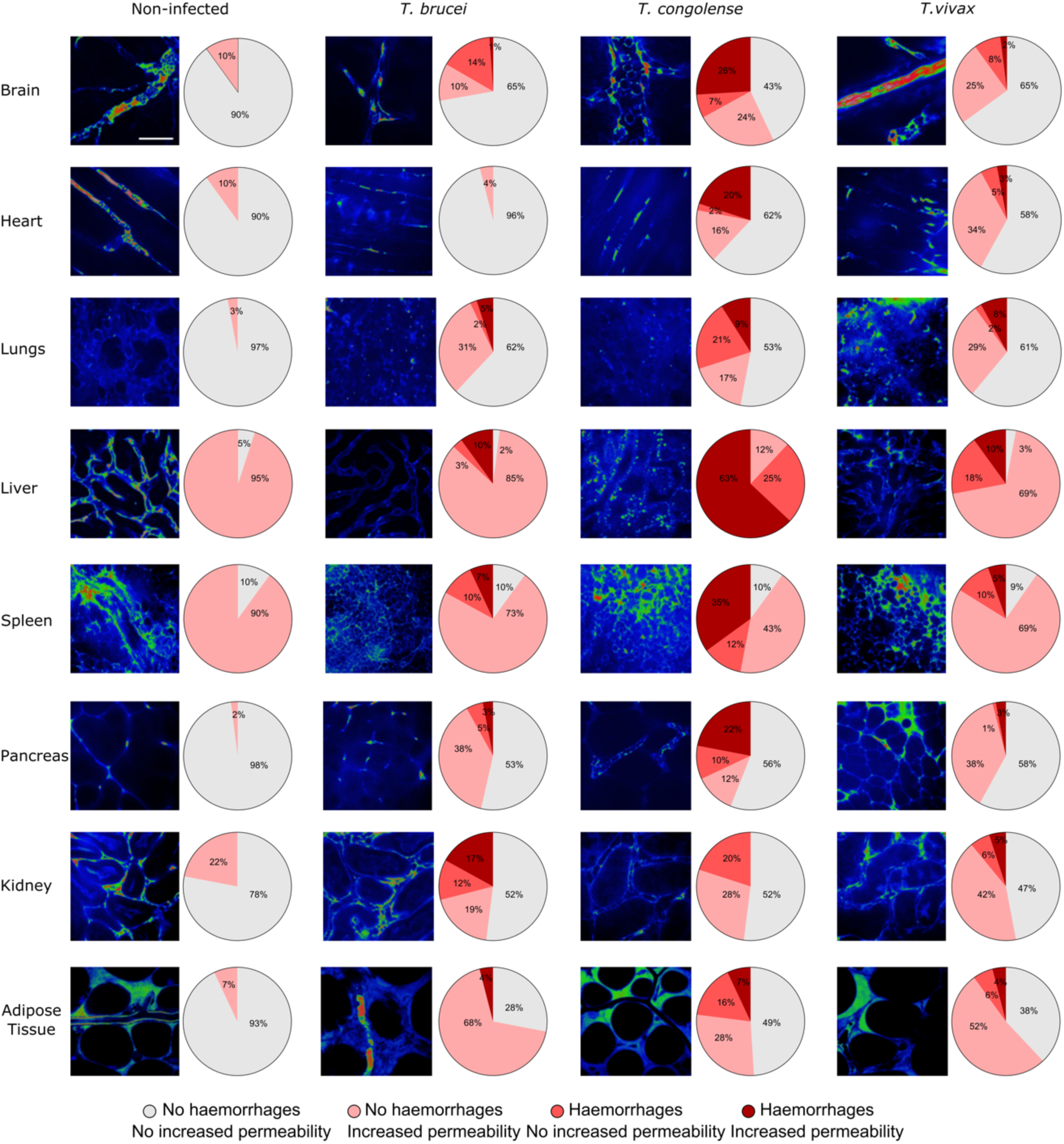
Prevalence of hemorrhagic lesions in C57B/6J mice infected with African trypanosomes, at the first peak of parasitaemia. For each organ, representative images and quantifications of the number of vessels with vascular damage by increased permeability, hemorrhagic lesions, or both, are shown. Scale bar is 40 µm. Heatmaps are color-coded according to key. Data from *T. brucei* and *T. congolense* (brain only) infections was re-analyzed from De Niz *et al*. 2021 and Silva Pereira *et al*. 2022, respectively.

### African trypanosomes induce different responses in the expression of key endothelium adhesion molecules

Having established that African trypanosomes cause distinct damage to blood vessels across multiple organs, we then analyzed the vascular endothelial cell response to trypanosome infection. Our previous work and others have shown that host-parasite interactions at the vascular endothelial interface are mediated by and/or result in up- or down-regulation of key vascular endothelial adhesion molecules such as ICAM1, ICAM2, CD36, E- and P-selectin, ESAM, VCAM1 and PECAM1. We selected the adhesion molecules to analyze based on their previously studied relevance to *Trypanosoma* infections (De Niz et al., 2021; Girard et al., 2005; Masocha et al., 2007; Silva Pereira et al., 2022) as well as other protozoan parasites (Bachmann et al., 2022; Cunningham et al., 2017; De Niz et al., 2016; Hopp et al., 2015; Ross et al., 2021). We used intravital imaging and fluorescence microscopy to quantify how the expression of key endothelial cell adhesion molecules changed upon infection, focusing on the first peak of parasitemia of each *Trypanosoma* species. **Figure 4A** shows representative images of the brain, for mice infected with each *Trypanosoma* species, focusing on each vascular adhesion molecule. Representative images of remaining organs are shown in **Supplementary Figures 1-8**.

**Figure 4.**
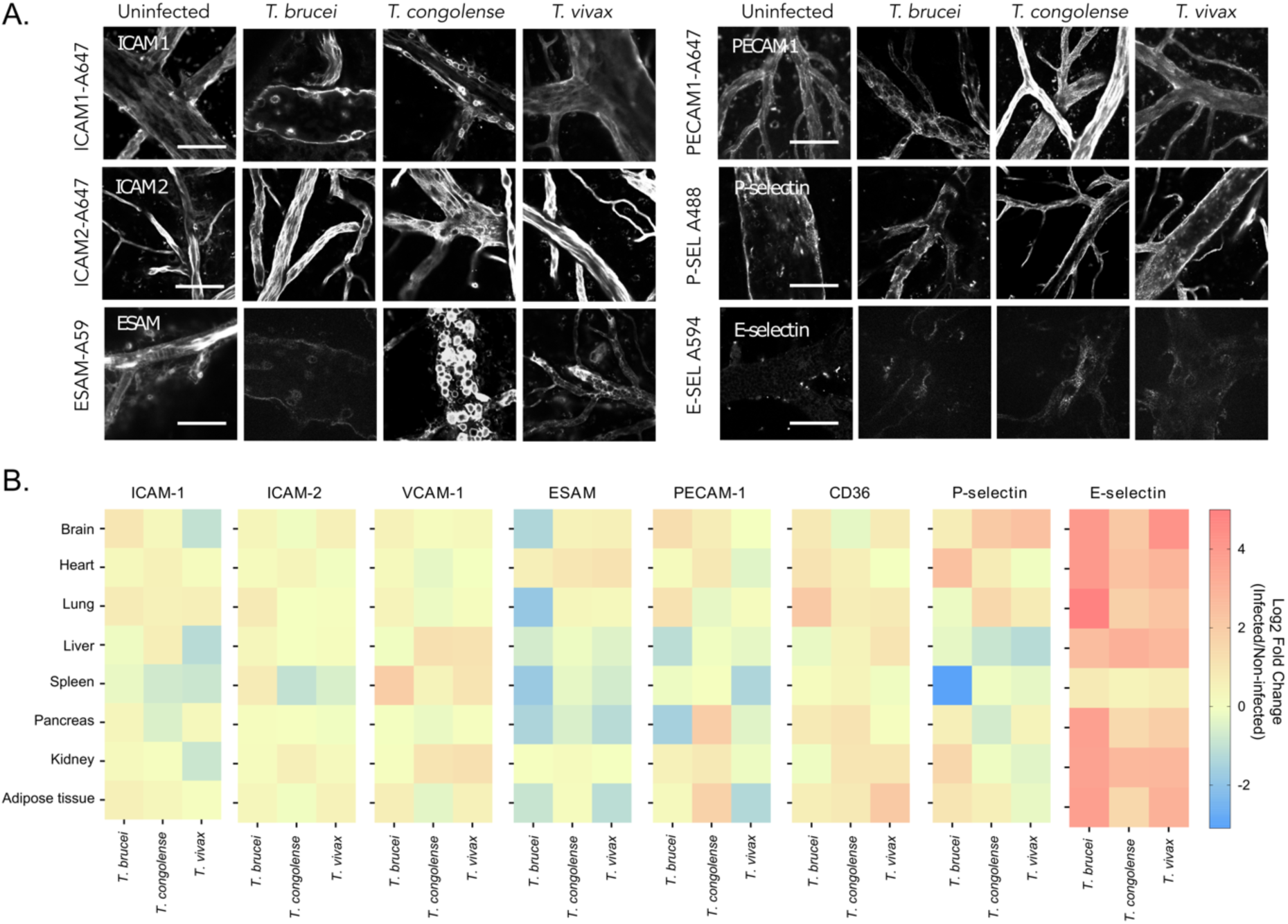
Expression of endothelial cell surface proteins at the first peak of parasitaemia of African trypanosome infections in C57B/6J mice. A. Representative images of each endothelial cell surface protein in the brain vasculature, detected by fluorescence microscopy. Scale bar is 40 µm. B. Heatmap of log_2_ fold change of protein (intercellular adhesion molecule-1, ICAM-1; intercellular adhesion molecule-2, ICAM-2; vascular cell adhesion molecule-1, VCAM-1; endothelial cell adhesion molecule, ESAM; Platelet endothelial cell adhesion molecule, PECAM-1; CD36; P-selectin; or E-selectin) expression relative to non-infected controls. Heatmaps are color-coded according to key. Data from *T. brucei* (all proteins, but ESAM) and *T. congolense* (ICAM1, ICAM2, VCAM-1) infections was reused from De Niz *et al*. 2021 and Silva Pereira *et al*. 2022, respectively.

We observed that the endothelial cell response significantly depended on both, the organotypic microenvironment, and the African trypanosome species. **Figure 4B** shows the relative change in fluorescence intensity in the vasculature of each organ upon infection with the three African Trypanosome species at the peak of parasitemia. This is expressed as a log2 fold-change relative to uninfected mice. ICAM-1 (CD54) (**Figure 4B, left-most panel**) encodes a cell surface glycoprotein typically expressed in endothelial cells and immune cells. Upon cytokine stimulation, its concentration can increase significantly. ICAM-1 plays an important role in cell-cell adhesion, leukocyte extravasation, modulation of inflammation, and host-pathogen interactions. We detected a significant decrease in ICAM-1 expression in the liver of *T. vivax*-infected mice (Log_2_ Fold Change (LFC)=−1.17). Representative images of ICAM-1 in organs where infection with any of the *Trypanosoma spp.* caused up-regulation greater than 1.5-fold or down-regulation under 0.5-fold compared to control, are shown in **Supplementary Figure 1**. Similar to ICAM-1, ICAM-2 (CD102) (**Figure 4B, second panel**) also plays a major role in mediation of inflammatory responses and immune cell trafficking, including cell-cell adhesion, lymphocyte recirculation and extravasation. ICAM-2 expression values did not change significantly throughout infection (−1 < LFC > 1). Representative images of ICAM-2 in organs where infection with any of the *Trypanosoma spp.* caused up-regulation greater than 1.5-fold or down-regulation under 0.5-fold compared to control, are shown in **Supplementary Figure 2**. VCAM-1 (CD106) (**Figure 4B, third panel**) also mediates adhesion of leukocytes to the vascular endothelium, and its upregulation is mediated by the presence of cytokines such as TNFa, IL1 and IL4. VCAM-1 expression was significantly increased in the spleen of *T. brucei*-infected mice (LFC=1.96), liver (LFC=1.21) and kidney (LFC=1.05) of *T. congolense*-infected mice, as well as liver (LFC=1.12), kidney (LFC=1.20), and spleen (LFC=1.06) of *T. vivax*-infected mice. Representative images of VCAM-1 in organs where infection with any of the *Trypanosoma spp.* caused up-regulation greater than 1.5-fold or down-regulation under 0.5-fold compared to control, are shown in **Supplementary Figure 3**. ESAM (endothelial cell adhesion molecule) (**Figure 4B, fourth panel**) is also involved in cell-cell adhesion, bicellular tight junction assembly, and regulation of actin cytoskeleton organization. ESAM was decreased in brain (LFC=−1.36), lungs (LFC=−1.85), spleen (LFC=−1.77), and pancreas (LFC=−1.40) of *T. brucei*-infected mice. It was also reduced in the adipose tissue (LFC=−1.13) and pancreas (LFC=−1.18) of *T. vivax*-infected mice, but elevated in the heart (LFC=1.13). Representative images of ESAM in organs where infection with any of the *Trypanosoma spp.* caused up-regulation greater than 1.5-fold or down-regulation under 0.5-fold compared to control, are shown in **Supplementary Figure 4**. PECAM-1 (CD31, platelet and endothelial cell adhesion molecule 1) (**Figure 4B, fifth panel**) makes up a large portion of endothelial cell intercellular junctions, and it is thought be involved in leukocyte migration, angiogenesis and integrin activation. PECAM-1 was elevated in the brain (LFC=1.27) and lungs (LFC=1.14) of *T. brucei*-infected mice, but reduced in the liver (LFC=−1.13) and pancreas (LFC=−1.54). Interestingly, PECAM-1 was increased in the adipose tissue (LFC=1.75) and pancreas (LFC=1.94) of *T. congolense*-infected mice, but reduced in the adipose tissue (LFC=−1.29) and spleen (LFC=−1.37) of *T. vivax*-infected mice. Representative images of PECAM-1 in organs where infection with any of the *Trypanosoma spp.* caused up-regulation greater than 1.5-fold or down-regulation under 0.5-fold compared to control, are shown in **Supplementary Figure 5**. CD36 (**Figure 4B, sixth panel**) is involved in lipid transport, regulation of metabolic processes, and regulation of reactive oxygen species. It has biased expression in the adipose tissue, heart, lungs and mammary gland. CD36 was high in the heart (LFC=1.09) and lungs (LFC=2.09) of *T. brucei*-infected animals, kidney (LFC=1.00) and pancreas (LFC=1.09) of *T. congolense*-infected animals, as well as liver (LFC=1.04) and adipose tissue (LFC=2.12) in *T. vivax* infections. Representative images of CD36 in organs where infection with any of the *Trypanosoma spp.* caused up-regulation greater than 1.5-fold or down-regulation under 0.5-fold compared to control, are shown in **Supplementary Figure 6**. P-selectin (**Figure 4B, seventh panel**) is a type 1 transmembrane protein that functions as a cell adhesion molecule on the surface of endothelial cells, and activated platelets. It is upregulated in the presence of inflammatory cytokines. P-selectin plays a key role in the initial recruitment of leukocytes to the site of inflammation, and in the recruitment and aggregation of platelets at sites of injury. P-selectin was elevated in the heart (LFC=2.44), kidney (LFC=1.54), and adipose tissue (LFC=1.01) during *T. brucei* infections, but reduced in the spleen (LFC=−3.02). In *T. congolense* infections, P-selectin was elevated in the brain (LFC=2.05) and lungs (LFC=1.52). In *T. vivax* infections, P-selectin was also elevated in the brain (LFC=2.41), but reduced in the liver (LFC=−1.20). Representative images of P-selectin in organs where infection with any of the *Trypanosoma spp.* caused up-regulation greater than 1.5-fold or down-regulation under 0.5-fold compared to control, are shown in **Supplementary Figure 7**. Finally, E-selectin (**Figure 4B, eight panel**) is expressed only on endothelial cells activated by cytokines. Like P-selectin, it mediates cell-tethering and rolling interactions of cells expressing E-selectin ligands (including a plethora of leukocytes). We observed an increase in E-selectin expression in the vasculature of all organs, with the sole exception of the spleen, in response to the three trypanosome species. Representative images of E-selectin in organs where infection with any of the *Trypanosoma spp.* caused up-regulation greater than 1.5-fold or down-regulation under 0.5-fold compared to control, are shown in **Supplementary Figure 8**. A detailed comparison of relative mean fluorescence units representing expression of each receptor, per organ, during each infection is shown in **Supplementary Figure 9**. From these results, we conclude that African trypanosomes induce different responses in the endothelium and that those responses are organ-dependent.

### *T. vivax* cytoadheres to the mouse vasculature in an organ-dependent manner

*T. congolense* and *T. brucei* display inherently different behaviours in the blood vasculature, with the former being exclusively intravascular and sequestering to the endothelium (Losos et al., 1971; Silva Pereira et al., 2022), whilst the latter traverses the endothelium to proliferate in the extravascular spaces of tissues (Capewell et al., 2016; De Niz et al., 2021; Mabille et al., 2022; Trindade et al., 2016). However, to our knowledge, there is no consensus on the behaviour of *T. vivax* parasites in the vasculature, i.e. whether they extravasate, sequester, or remain intravascular but non-sequestered (discussed in Silva Pereira et al., 2019). Therefore, we questioned whether the obvious differences in endothelial cell adhesion molecule expression could be linked to the parasite interaction with the endothelium. To test this hypothesis, we investigated the behaviour of *T. vivax* parasites in the vasculature by intravital microscopy. We did not find evidence for *T. vivax* extravasation. On the contrary, we found that *T. vivax* parasites adhere to the vasculature in a process that resembles *T. congolense* sequestration, albeit with distinct vessel and tissue distributions.

*T. vivax* parasites sequester to the endothelium of all major organs (**Figure 5A-H**). The liver is the organ where parasites accumulate first, at day 1 p. i., mostly in their sequestered form (**Figure 5D**), although they find their way to the vasculature of remaining organs from day 2 p. i.. With some small fluctuations, the number of sequestered and non-sequestered parasites throughout infection increases in all organs but the liver. Here, parasites accumulate greatly during the first 2 days of infection. Then, gradually, parasite load reduces until day 6 p. i., after which it rises again. Furthermore, we observed that sequestration generally correlates linearly with parasite load of each organ’s vasculature (r^2^=0.98 for all organs, except lungs (r^2^=0.90), Pearson’s correlation). In fact, the vast majority of parasites exists sequestered to the endothelium throughout infection, with the sole exception to this behaviour being observed in the lungs (**Figure 5I**). Here, after day 4 post-infection, there is a significant reduction in the percentage of sequestered parasites, even though the number of free (i.e. non-sequestered) parasites gradually increases (q-value= 0.004, multiple unpaired t-test with False Discovery Rate approach) (**Figure 5C and 5I**). Sequestered parasites were distinguished from non-sequestered parasites by observation of their motility in videos captured by IVM (**Supplementary Videos 1-5**). The intravascular environment is visible due to the administration of FITC-Dextran, whereas parasites are identified by nuclei staining with Hoechst dye (**Figure 5J**). Upon closer observation, we detected that most parasites appear attached to the endothelium by their flagellum (**Figure 6A**), consistent with what has been previously described for *T. congolense* (Hemphill and Ross, 1995). To confirm our results by an alternative method, we compared the quantifications obtained by IVM with quantifications by qPCR following mouse perfusion and observed a high positive correlation (Pearson’s R^2^ = 0.64, p-value = 0.01) (**Figure 6B**). Perfusion immediately post-euthanasia removes non-sequestered parasites from circulation, which allows estimation of sequestered parasite load in individual organs.

**Figure 5.**
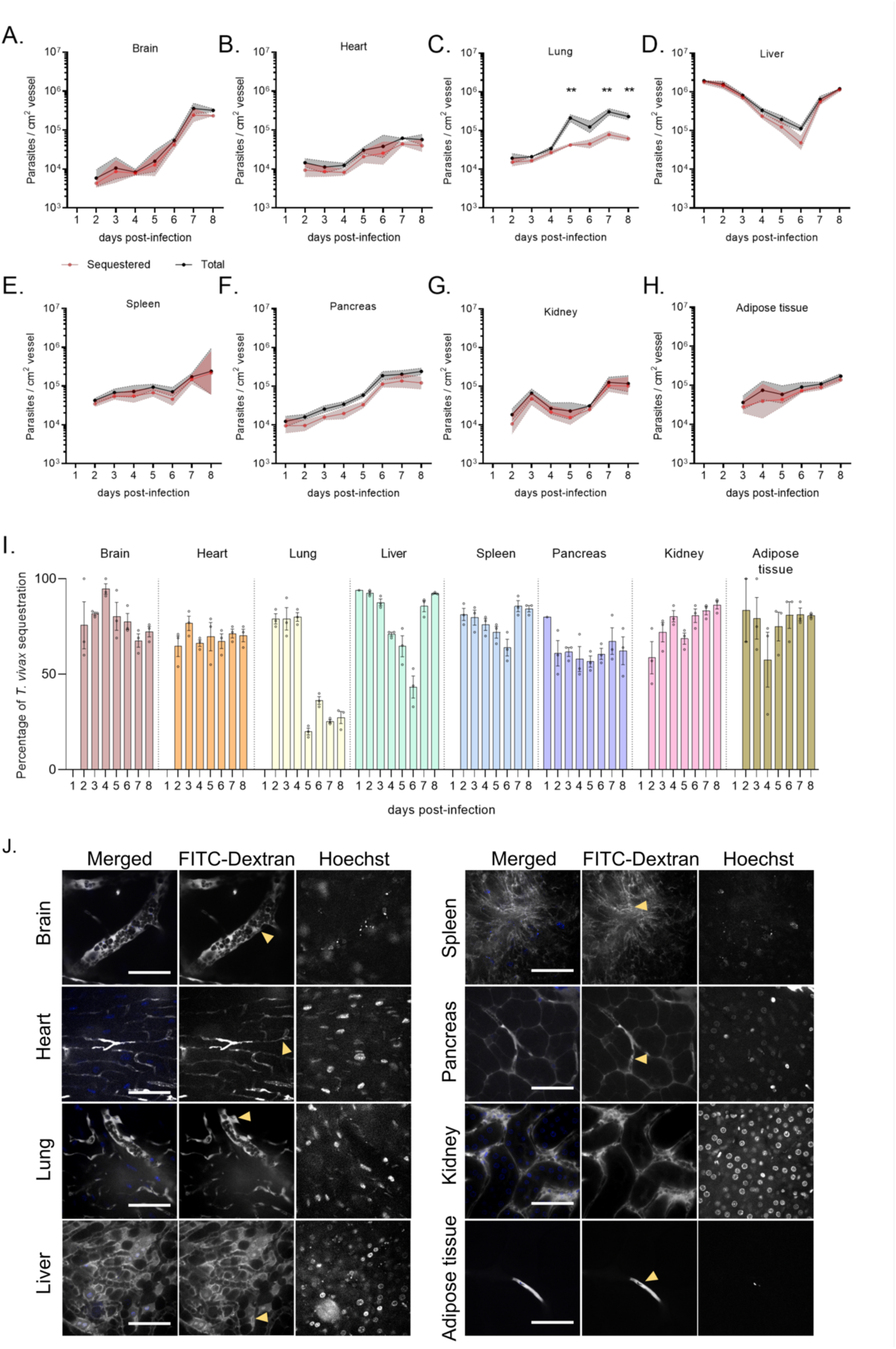
Dynamics of *T. vivax* sequestration during the first peak of parasitaemia. A-H. *T. vivax* total parasite load (black) and sequestered parasite load (red) throughout the first peak of parasitaemia in major organs. I. Percentage of *T. vivax* sequestration in major organs calculated as the number of sequestered parasites per cm^2^ of vessel divided by the total number of parasites detected per cm^2^ of vessel. Mean±SEM. J. Representative images of *T. vivax* sequestering to endothelial cells *in vivo*. Parasites are indicated with yellow arrow heads, intravascular environment is visible with FITC-Dextran, nuclei are stained with Hoechst. Scale bar is 40 µm.

**Figure 6.**
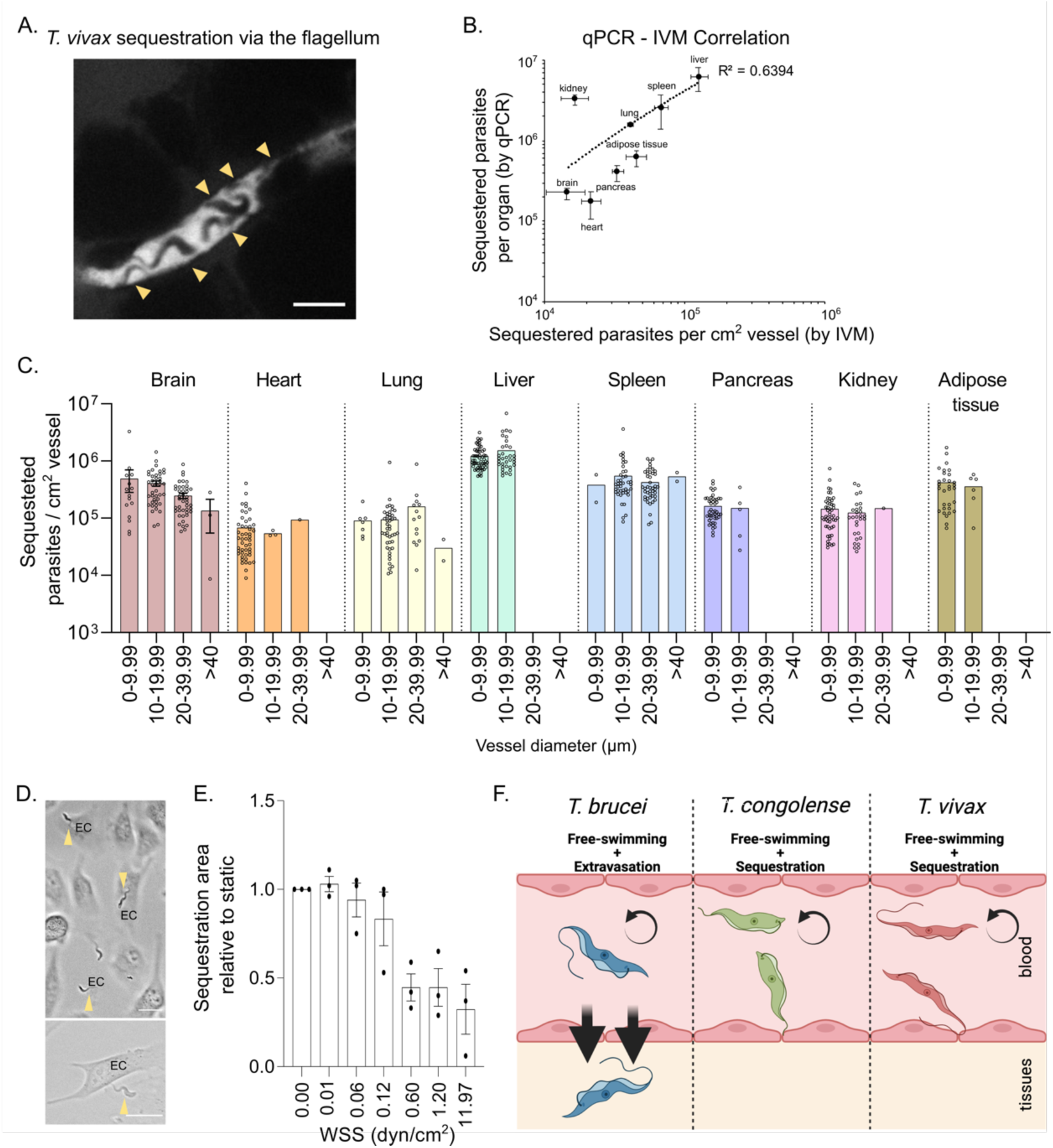
Distribution and reproducibility of *T. vivax* sequestration. A. Representative image of *T. vivax* sequestration, suggesting adhesion by the flagellum. Parasites are indicated with yellow arrow heads. Scale bar is 20 µm. B. Correlation between sequestered parasite load estimated by quantitative PCR (18S DNA) and by intravital microscopy. R^2^ = 0.6394, *Pearson’s* correlation, *p*-value < 0.001. C. *T. vivax* parasite distribution per vessel caliber in major organs. D. *T. vivax* adhesion ability at increasing wall shear stresses, compared to adhesion in static conditions (no flow). Mean ± SEM. E. Model of African trypanosome (*T. brucei*, *T. congolense*, and *T. vivax*) interaction with endothelial cells. All three species of parasites exist as free-swimming organisms in the blood. However, whilst *T. brucei* can extravasate the endothelium towards extravascular spaces of tissues, *T. congolense* and *T. vivax* remain adhered to the vasculature, in a process named sequestration.

We then investigated how parasites distributed by vessel diameter in each organ, which could reveal a potential preference for particular types of vessels. We divided all vessels imaged during day 8 p.i. (i.e. the peak of parasitemia for *T. vivax*) into four categories based on their diameter: 0-9.99µm, 10-19.99µm, 20-39.99µm, and those larger than 40µm. We then quantified the number of sequestered parasites present in each vessel. It is worth noting that, by IVM, the accessibility to large vessels is limited in some organs, such as the heart, the liver, the pancreas, or the adipose tissue. Therefore, in some organs, we did not capture vessels of larger diameters. Nonetheless, with the data we compiled, we did not detect any bias in sequestration towards particular vessel diameters. This is supported by the fact that the average number of sequestered parasites per cm^2^ vessel is rather constant in each organ, regardless of the diameter of the vessel (**Figure 6C**).

Finally, we tested whether the sequestration behavior could be mimicked *in vitro* and assessed the strength of the parasite-endothelium interaction. To achieve this, we pre-coated a 6 channel μ-Slide with bovine aorta endothelial cells (BAOECs), added *T. vivax* bloodstream form parasites previously extracted from mouse blood, and let them interact for 1 hour. After this time, we perfused the channels and counted the number of parasites in the same sample over increasing levels of flow rates, which correspond to increasing wall shear stress values. **Figure 6D** shows *T. vivax* parasites sequestered to BAOECs after sustaining wall shear stress of 0.12 dyn/cm^2^. We observed no change in parasite numbers until a wall shear stress of 0.6 dyn/cm^2^ is applied. At this force, approximately half of the parasites detach, whilst the other half remains sequestered as the stress increases to close to 12 dyn/cm^2^. These wall shear stress values are consistent with the arteriole-capillary-venule unit and suggest that the interaction between *T. vivax* and the endothelial cell is strong and active. These results confirm that *T. vivax* parasites sequester to the vascular endothelium both *in vivo* and *in vitro*, with 40% of the parasite population being able to sustain a wall shear stress of at least 12 dyn/cm^2^ (**Figure 6E**).

## Discussion

Gaining insight into parasite-host vascular interactions in the context of *Trypanosoma* infections is pivotal for a) our understanding of clinically-relevant pathology induced by the various *Trypanosoma* species; b) identifying suitable animal models to replicate veterinary and human-relevant symptoms and syndromes; and c) the development of therapeutic strategies for the treatment and control of *Trypanosoma spp.-*induced pathology, both during acute and chronic infection, and post-infection resolution.

In our work, we focused on the comparison of survival, vascular pathology, and sequestration patterns (or lack thereof) of 3 *Trypanosoma* species: *T. brucei, T. congolense,* and *T. vivax.* We began by showing major differences in host survival upon infection with specific strains of each species. Interestingly, the longest survival is displayed by hosts infected with *T. brucei,* a species that succeeds in traversing the host vascular endothelium to establish extravascular parasite reservoirs. Meanwhile, *T. congolense* and *T. vivax* display no extravasation. Upon comparing vascular permeability changes induced by *Trypanosoma spp.* interactions, we observed that *T. brucei* causes the least severe effects on vascular homeostasis. Conversely, *T. congolense* and *T. vivax* infections result in significant vascular pathology, including extensive microhemorrhages, which likely contribute to the severe organ-specific pathology of the diseases they cause. We have discussed our findings on *T. brucei* and *T. congolense* extensively in our previous work (De Niz et al., 2021; Silva Pereira et al., 2022), while our findings on *T. vivax* are presented for the first time. The organs most affected by each of the two non-extravasating species are widely different: while *T. congolense* causes the most severe effects in the liver and spleen, *T. vivax* does so in the heart and adipose tissue, despite sequestration being high across most organs. Of particular interest relevant to both *T. congolense* and *T. vivax* is the sudden vascular pathology observed in the lungs. Given the lung’s role in blood oxygenation and the volume of blood this organ receives, we hypothesize that this is the cumulative result of parasite circulation, adhesion, and the circulation of inflammatory mediators that result in lung mediated damage, similar to what has been described for *Plasmodium* infections.

*T. vivax* leads to a lower profile of microhemorrhages across all organs, compared to *T. congolense.* The different phenotype between these two species raises important questions regarding the mechanisms of parasite-host interactions that result in such severe vascular changes in an organ- and species-specific manner. For instance, how are vascular permeability and microhemorrhages induced? Based on work on these and other protozoan parasites, we suggest that various possibilities exist, including direct interactions by the flagellum with the endothelium enabling probing and adhesion; extracellular vesicle release; and immunopathological responses including inflammatory cytokine release and induction of immune cell recruitment. We explored here whether a correlation exists between sequestered parasites and vascular permeability at day 8 p.i., but we didn’t find one (r^2^=0.06, *p*-value = 0.55) suggesting that although sequestration is an important factor, it is not the main factor mediating *T. vivax* effects on vascular homeostasis. We envisage our future work will explore other key parameters such as cytokine profiling, extravascular vesicle release, direct binding of *T. vivax* and *T. congolense*, and flagellar probing and dynamics at the vascular endothelium.

One aspect we explored in our work, potentially linked to both, vascular permeability and sequestration, is the up- and down-regulation of receptors at the vascular endothelium. This alone has been a topic of great interest across parasitology fields. Amongst the best studied parasites in terms of ligand-receptor interactions is *Plasmodium*, which has evolved a complex export machinery dedicated to transporting virulence factors to the surface of the red blood cell to enable parasite sequestration (Boddey and Cowman, 2013; Maier et al., 2002; Przyborski et al., 2003). A vast amount of the *Plasmodium* genome is dedicated to generate *var* gene repertoires (reviewed in Deitsch and Hviid, 2004; Scherf et al., 2008; Smith et al., 2001) that enable organ-specific ligand-receptor interactions with the vascular endothelium of the mammalian host, while ensuring immune evasion. Many host receptors have been well-characterized in terms of their relevance for *Plasmodium* sequestration (reviewed in Hviid and Jensen, 2015). Conversely, relatively little is known about ligand-receptor interactions relevant to *Trypanosoma spp.* and their mammalian hosts, with our group recently exploring the relevance of ICAM1, ICAM2, PECAM1, VCAM, CD36, ESAM, and P- and E-selectin for *T. brucei* extravasation (De Niz et al., 2021) and *T. congolense* sequestration in the brain (Silva Pereira et al., 2022). Equally, little is known about the relevance of potential ligands at the surface of *Trypanosoma spp.* mediating these interactions, and/or whether variant surface glycoproteins (VSGs), analogous to *var* genes, play a role in this phenomenon. In our current work, we showed that the overall profile of receptor expression is significantly different between *Trypanosoma spp.,* with *T. vivax* causing most significant changes in ICAM1, VCAM1 and E-selectin. Important further work will include exploring the effect of receptor abrogation (by knock-out or antibody blocking) on *T. vivax* distribution and pathology. Similarly, that these receptors are significantly changed hints towards possible *T. vivax* candidates that could act as ligands.

Besides triggering different responses from the vascular endothelium, African trypanosomes display inherently different behaviors in the blood in terms of their motility, which tissues they accumulate in, and how they physically interact with the endothelium. Whilst the first aspect has been described in detail elsewhere (Bargul et al., 2016), here we presented evidence that *T. vivax* sequesters to the vascular endothelium and described its tissue distribution in a mouse model.

The majority of *T. vivax* parasites in the mouse blood exists in their sequestered form, which was also observed for *T. congolense* (Silva Pereira et al., 2022). This indicates that sequestration is advantageous for these parasites. Although the reasons behind it remain unclear, we postulate that sequestration enhances parasite-endothelium interactions by facilitating the hijack of cellular functions and/or host’s nutrients. For instance, sequestration might aid iron uptake, which is both essential for trypanosome survival and a source of host pathology due to anemia (Stijlemans et al., 2015). This is consistent with earlier results showing that *T. congolense* survives in co-culture and sequesters to both fixed and live BAOECs, but only multiplies if endothelial cells are alive (Hemphill et al., 1994). Alternatively, sequestration may help immune evasion by preventing splenic clearance, as observed in malaria (Buffet et al., 2011; del Portillo et al., 2012; Ghosh and Stumhofer, 2021).

Unlike *T. brucei*, where there is a clear preference for particular organs, such as the pancreas (De Niz et al., 2021) and the adipose tissue (Trindade et al., 2016), we did not detect any obvious tropism of *T. vivax*. In contrast, we saw a steady increase in both total and sequestered parasite load in all organs with the exception of the liver, perhaps due to its role in parasite clearance, as observed for *T. cruzi* (Sardinha et al., 2010). This also differs from *T. congolense*, where despite presence of parasites in the vasculature of all organs, there is a strain-specific preference for particular tissues, directly linked to pathology (Silva Pereira et al., 2022).

A final thing that transpires from this work is that sequestration might be a trait older than current African trypanosome species. Whilst sequestration has for long been assumed to be *T. congolense*-specific, our data clearly shows that *T. vivax* also employs it as a survival mechanism. Yet, whether this is a species-wide feature or specific to some strains, remains unknown. *T. vivax* split from the last common ancestor before *T. brucei* and *T. congolense*. Since both *T. vivax* and *T. congolense* sequester, but not *T. brucei* (Fig. 6F), either *T. brucei* has lost the ability to do it, or sequestration was independently acquired by *T. congolense* and *T. vivax*. The first hypothesis is undoubtedly more parsimonious, (i.e. requiring the occurrence of fewer independent events), and would indicate that sequestration is an ancient trait carried over from the last common ancestor of African trypanosomes. Our future work will aim to investigate the evolutionary dynamics of genes involved in sequestration, with the view to assess their origin. Moreover, given the marked genetic differences between T. vivax from East Africa, South Africa, and West Africa/South America (Cortez et al., 2006; Rodrigues et al., 2008, 2017; Silva Pereira et al., 2020), as well as the existence of *T. vivax*-like organisms (Rodrigues et al., 2008), it is important to compare and contrast how multiple strains behave in the bloodstream, before considering sequestration a species-wide trait. Ultimately, why *T. brucei* extravasates whilst *T. congolense* and *T. vivax* sequester is puzzling. Answering this question might help us understand the differences in pathology, disease severity and general parasite behavior observed amongst these three close relatives.

Several additional questions remain unanswered, including how are *T. congolense* and *T. vivax* parasites binding to endothelial cells at the molecular level, how does endothelial cell activation affect sequestration, and ultimately, whether preventing sequestration could abrogate disease. With this work, we lay the ground for answering these important questions by presenting sequestration as a previously unidentified behavior of *T. vivax* and describing several candidates for host ligands of sequestration.

## Materials and Methods

### Animal experiments

This study was conducted in accordance with EU regulations and ethical approval was obtained from the Animal Ethics Committee of Instituto de Medicina Molecular (AWB_2016_07_LF_Tropism and AWB_2021_11_LF_TrypColonization). Infections were performed at the rodent facility of Instituto de Medicina Molecular, in 6–10 weeks old, wild-type, male C57BL/6 J mice (Charles River, France). Mice were infected by intraperitoneal injection (i. p.) of 2 × 10^3^ *T. congolense* savannah 1/148 (MBOI/NG/60/1–148) (Young and Godfrey, 1983), 2 × 10^4^ *T. brucei* TY1-TdTomato-FLuc AnTat1.1E (Calvo-Alvarez et al., 2018), or 1 × 10^4^ *T. vivax* Y486 (Gibson, 2012). Parasitemia was estimated daily by hemocytometry from tail venipuncture. Mice were sacrificed by anesthetic overdose or CO2 narcosis. Blood was collected by heart puncture and, when necessary, mice were perfused with 50 mL heparinized PBS. For molecular biology, tissues were dissected, washed in PBS and immediately imaged or snap frozen in liquid nitrogen.

### Intravital and *ex vivo* imaging

For intravital imaging, surgeries were separately performed in the brain; the lungs and heart; the liver; the pancreas, spleen and kidney; the adipose tissues and lymph nodes, as described in De Niz and Figueiredo (2022). Briefly, mice were anaesthetized with a mixture of ketamine (120 mg/kg) and xylazine (16 mg/kg) injected intraperitonially. After checking for reflex responses and ensuring none occurred, mice were then intraocularly injected with Hoechst 33342 (stock diluted in dH_2_O at 100 mg/ml; injection of 40 μg/kg mouse), 70 kDa FITC-Dextran (stock diluted in 1x PBS at stock concentration of 100 mg/ml; injection of 500 mg/kg), and vascular markers (20 µg) of interest conjugated to AF647 (CD31, ICAM1, ICAM2, and CD36 (BioLegend), P-selectin (BD Pharmingen)), FITC or AF488 ((VCAM-1 (Invitrogen)), E-Selectin (BD PharMingen)) or AF594 (ESAM (BioLegend)). A temporary glass window (Merk rectangular coverglass #1.5, 100 mm x 60 mm or circular coverglass (12 mm)) was implanted in each organ, and secured either surgically, with surgical glue, or via a vacuum.

For intravital microscopy, all imaging relative to parasite quantifications, vascular density and vascular leakage was done in a Zeiss Cell Observer SD (spinning disc confocal) microscope (Carl Zeiss Microimaging, equipped with a Yokogawa CSU-X1 confocal scanner, an Evolve 512 EMCCD camera and a Hamamatsu ORCA-flash 4.0 VS camera) or in a 3i Marianas SDC (spinning disc confocal) microscope (Intelligent Imaging Innovations, equipped with a Yokogawa CSU-X1 confocal scanner and a Photometrics Evolve 512 EMCCD camera). Laser units 405, 488, 561 and 640 nm were used to image Hoechst in nuclei, extravascular and intravascular FITC-Dextran, TdTomato in *T. brucei,* and vascular markers respectively. The objective used to image vascular density, vascular leakage, and proportion of intravascular and extravascular parasites was a 40x LD C-Apochoromat corrected, water immersion objective with 1.1 NA and 0.62 WD. Twenty images were obtained in any one time lapse, with an acquisition rate of 5 frames per second. For all acquisitions, the software used was ZEN blue edition v.2.6 (for the Zeiss Cell Observed SD) allowing export of images in .czi format, and 3i Slidebook reader v.6.0.22 (for the 3i Marianas SD), allowing export of images in TIFF format.

### Vascular permeability quantification

In order to quantify vascular permeability changes, we took as reference the marker 70 kDa FITC-Dextran as previously published methodology (Egawa et al., 2013) and as previously described in our work (De Niz et al., 2021; Silva-Pereira et al., 2022). We measured FITC-Dextran in intravascular and extravascular regions in uninfected mice, and then at each time post-infection with each of the *Trypanosoma spp.* The permeability ratio was calculated using the following equation: 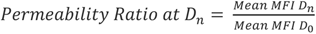, where mean MFI_Dn_ is the extravascular MFI at a specific Day *n*, and the mean MFI_D0_ is the extravascular MFI in uninfected mice (Day 0). We performed vascular permeability measurements in a minimum of 60 fields of view per day post-infection, per *Trypanosoma spp* in a minimum of 3 mice.

### Vascular density and diameter quantification

In order to quantify vascular density, we took as reference the vascular marker CD31-A647 and 70 kDa FITC-Dextran as previously described (De Niz and Figueiredo, 2022). Briefly, we acquired 100 fields of view, and for each field of view, the total area was defined as 100%. Using the CD31 signal we were able to segment out the vascular regions using Fiji. We calculated the percentage of vascular area covered using the following formula: 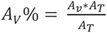 where A_T_ is the total area of the field of view, A_T%_ is 100, and A_v_ is the total area marked by CD31. Vessel diameters were measured using Fiji. Parasite density is expressed as number of parasites per cm^2^ of vessel. A minimum of 60 fields of view were quantified per days post-infection and *Trypanosoma spp*.

### Quantitative PCR

Tissues were homogenized mechanically in 500µl lysis buffer using silica beads and a tissue disruptor machine. Genomic DNA was extracted from a mass of homogenate equivalent to 25mg of solid tissue using NZYTech gDNA isolation kit. DNA was quantified by Spectrophotometry in the Nanodrop 2000 (Thermo Fischer Scientific). *T. vivax* 18S ribosomal DNA genes were amplified from genomic DNA and converted into parasite number using a standard curve.

### Detachment assay

Parasites were isolated from mouse by anion exchange chromatography (Lanham and Godfrey, 1970) and stained with 5 mM Vybrant CFDA SE Cell Tracer dye (#V12883, Invitrogen) diluted 1000 times in trypanosome dilution buffer (TDB) (5 mM KCl, 80 mM NaCl, 1 mM MgSO4, 20 mM Na2HPO4, 2 mM NaH2PO4, 20 mM glucose, pH 7.4), and incubated for 25 min at 34 °C, 5% CO2. At the end of the incubation period, parasites were washed and resuspended in TDB, added to the endothelial cell monolayers, and incubated for 1 hr at 34 °C, 5% CO_2_. Flow was applied with a perfusion syringe pump containing PBS at defined flow rates, for 1 minute each. Parasites were imaged live on a Zeiss LSM 980 (Carl Zeiss Microimaging) with a 20 X water-immersion objective (0.8 numerical aperture and 0.55mm working distance) before and after each flow session. We acquired 10 fields of view per condition (each WSS value), per replicate (3 replicates), with green laser (488nm, maximum power of 13mW). For all acquisitions, the software used was ZEN blue edition v.2.6, allowing export of images in czi format.

### Quantification and analysis

Data were displayed in graphs and heatmaps generated using Prism 9 software (GraphPad) and RStudio. For IVM and ex-vivo microscopy analyses, values were calculated from triplicate experiments with 3 biological replicates each, and/or at least 100 images per condition. Pearson correlations measures (R), and R^2^ values were calculated to determine the strength of linear association between parasite density and either vascular permeability or vascular density. For the analysis of the results from the detachment assay, we used Fiji software to count the area containing adhered parasites per field of view and generated graphs using Prism 9 software (GraphPad). Statistical details of experiments are included in the figure legends and the results section.

## Acknowledgements

We thank Dr Álvaro Acosta-Serrano (University of Notre Dame), Dr Brice Rotureau (Institut Pasteur), and Dr Loïc Rivière (University of Bordeaux) for providing *T. congolense* 1/148, triple reporter (TY1-TdTomato-FLuc) AnTat1.1E *T. brucei*, and *T. vivax* Y486 parasite lines, respectively. We thank Prof. Luisa M Figueiredo for guidance and support during our time in her laboratory, and for carefully proofreading this manuscript. We thank all members of the Figueiredo lab for their input and helpful discussions. We also thank Prof. Cláudio A Franco and his group for valuable scientific input, and for carefully proofreading this manuscript. Finally, we acknowledge the Bioimaging and Rodent facilities at iMM-JLA, and are especially grateful for the support of Jose Rino, Antonio Temudo, Iolanda Sousa Moreira, and Daniel Costa.

S.S.P. was funded by European Union’s Horizon 2020 research and innovation program through a Marie Skłodowska-Curie Individual Standard European Fellowship, under grant agreement no. 839960. This project received the support of a fellowship from “la Caixa” Foundation (ID 10001043). This work was supported by HFSP (LT000047/2019-L) and EMBO (ALTF 1048-2016) Individual Fellowships to M.D.N. All work was performed in the lab of Luisa M. Figueiredo. L.M.F. is an Investigator CEEC of the Fundação para a Ciência e a Tecnologia (CEECIND/03322/2108). This project has received funding from the European Research Council (ERC) under the European Union’s Horizon 2020 research and innovation programme (grant agreement No 771714).

