## Supplementary Figures for "Investigation of *Trypanosoma-*induced vascular damage sheds insights into *Trypanosoma vivax* sequestration"

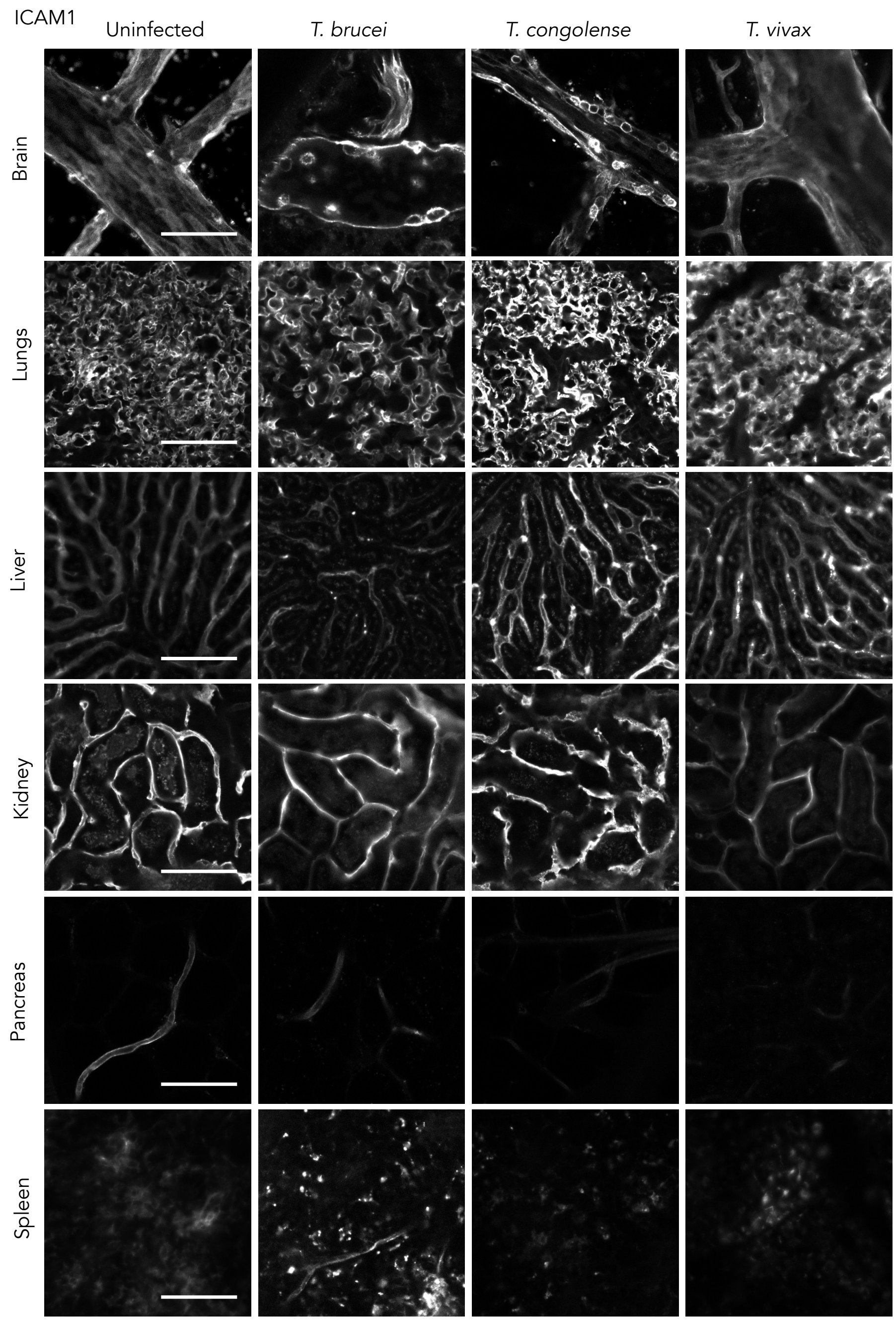


**Supplementary Figure 1. Expression ICAM-1 at the first peak of parasitaemia of African trypanosome infections in C57B/6J mice.** Representative images of ICAM-1 in the brain, lungs, liver, kidney, pancreas and spleen, detected by fluorescence microscopy. Scale bar is 40 μm.

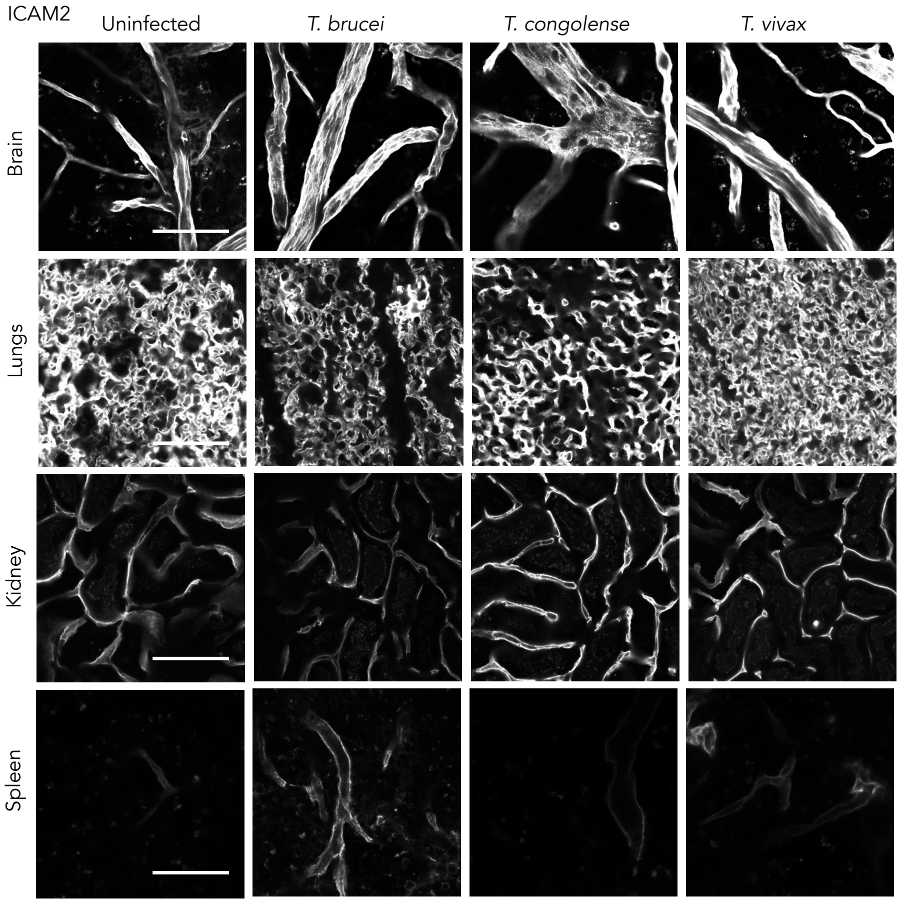


**Supplementary Figure 2. Expression ICAM-2 at the first peak of parasitaemia of African trypanosome infections in C57B/6J mice.** Representative images of ICAM-2 in the brain, lungs, kidney and spleen, detected by fluorescence microscopy. Scale bar is 40 µm.

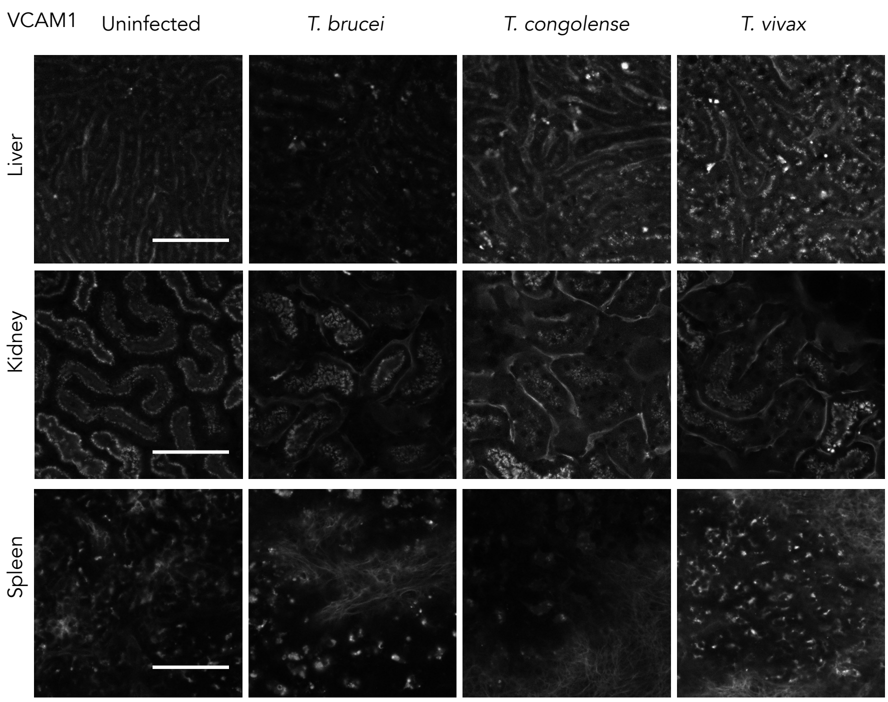


**Supplementary Figure 3. Expression VCAM-1 at the first peak of parasitaemia of African trypanosome infections in C57B/6J mice.** Representative images of VCAM-1 in the liver, kidney and spleen, detected by fluorescence microscopy. Scale bar is 40 μm.

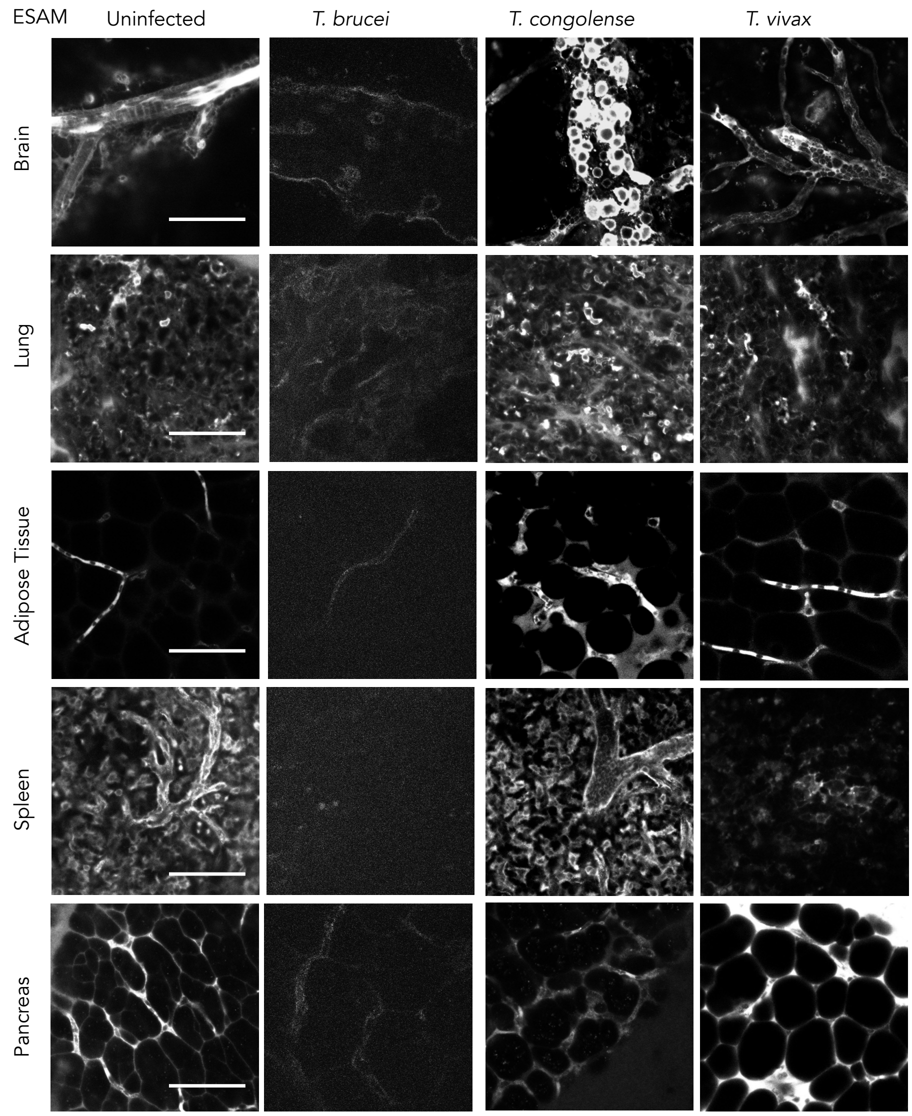


**Supplementary Figure 4. Expression ESAM at the first peak of parasitaemia of African trypanosome infections in C57B/6J mice.** Representative images of ESAM in the brain, lungs, adipose tissue, spleen and pancreas, detected by fluorescence microscopy. Scale bar is 40 μm.

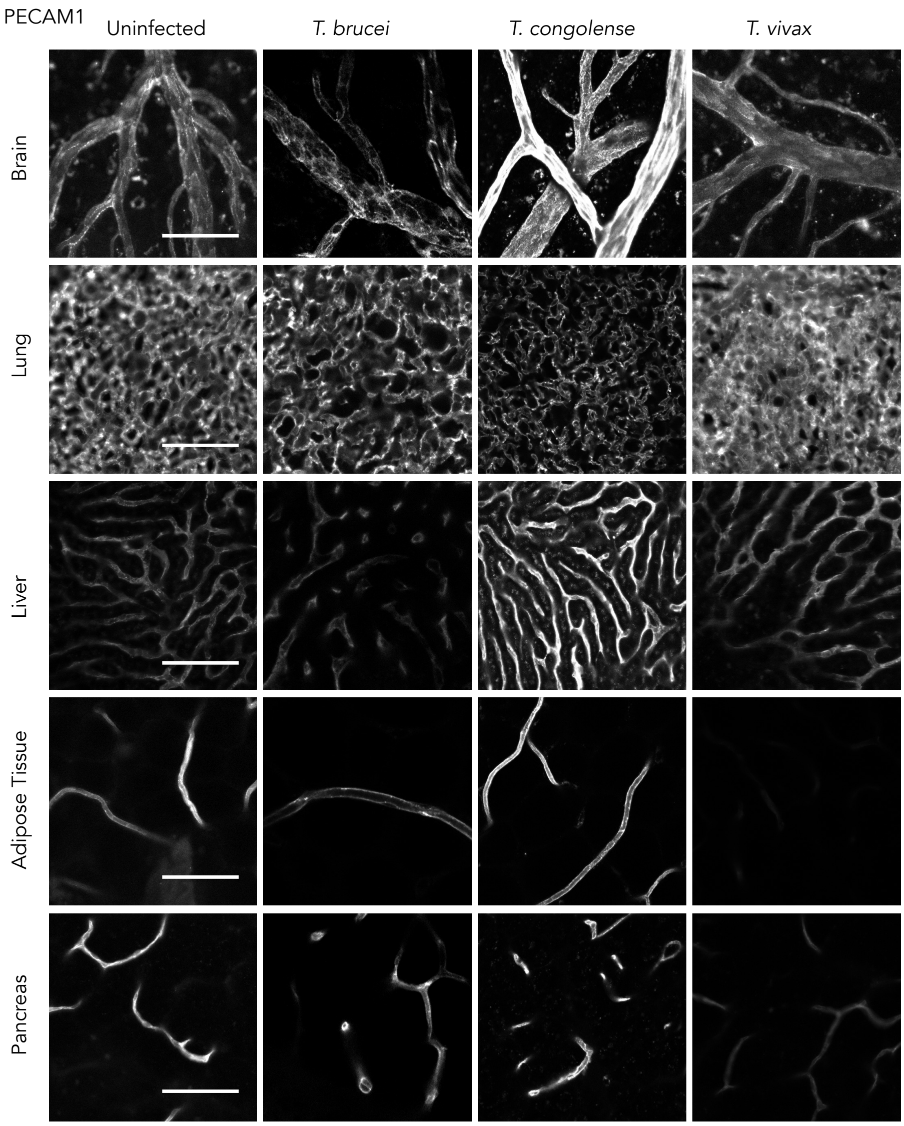


**Supplementary Figure 5. Expression PECAM-1 at the first peak of parasitaemia of African trypanosome infections in C57B/6J mice.** Representative images of PECAM-1 in the brain, lungs, liver, adipose tissue and pancreas, detected by fluorescence microscopy. Scale bar is 40 μm.

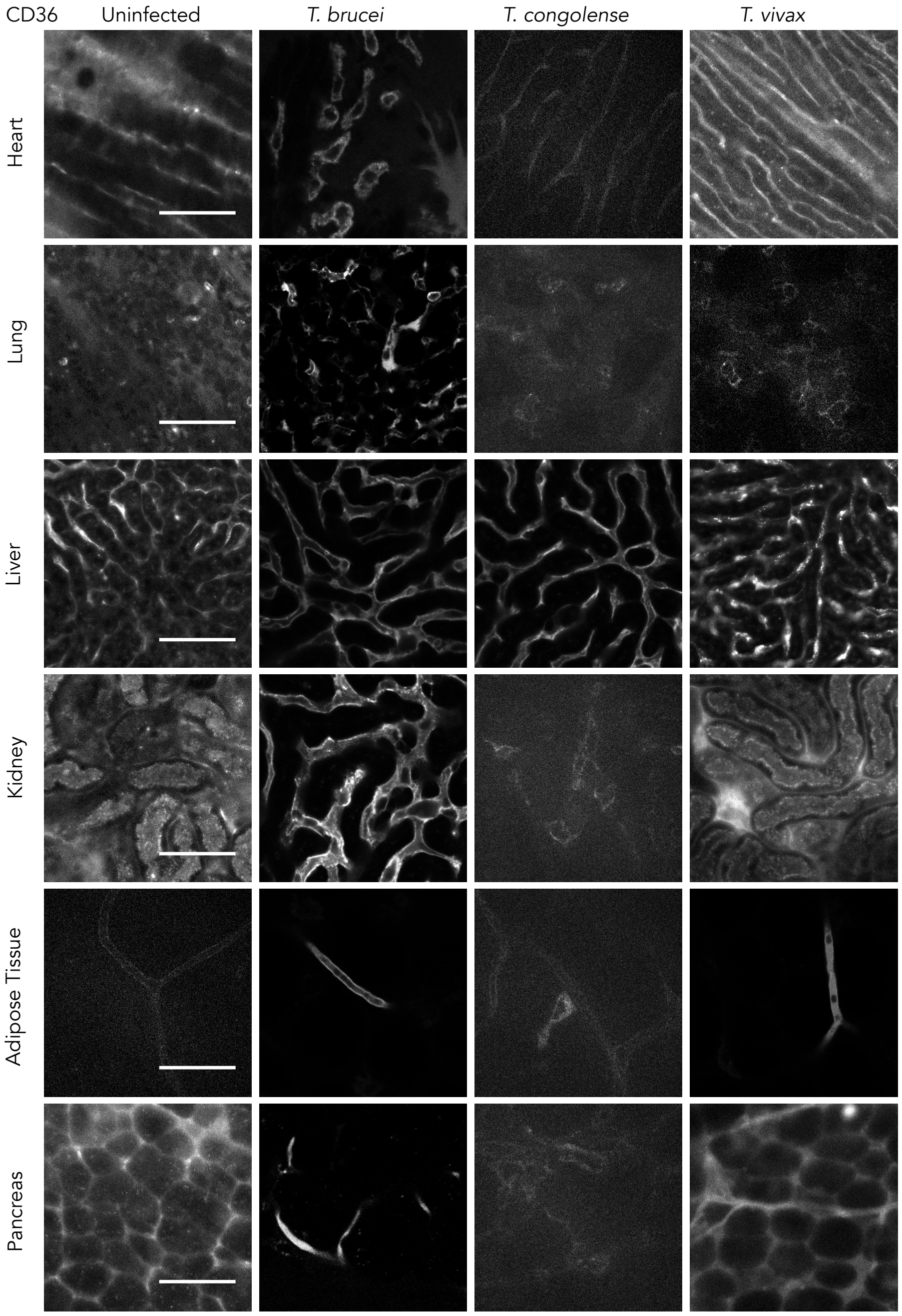


**Supplementary Figure 6. Expression CD36 at the first peak of parasitaemia of African trypanosome infections in C57B/6J mice.** Representative images of CD36 in the heart, lungs, liver, kidney, adipose tissue and pancreas, detected by fluorescence microscopy. Scale bar is 40 μm.

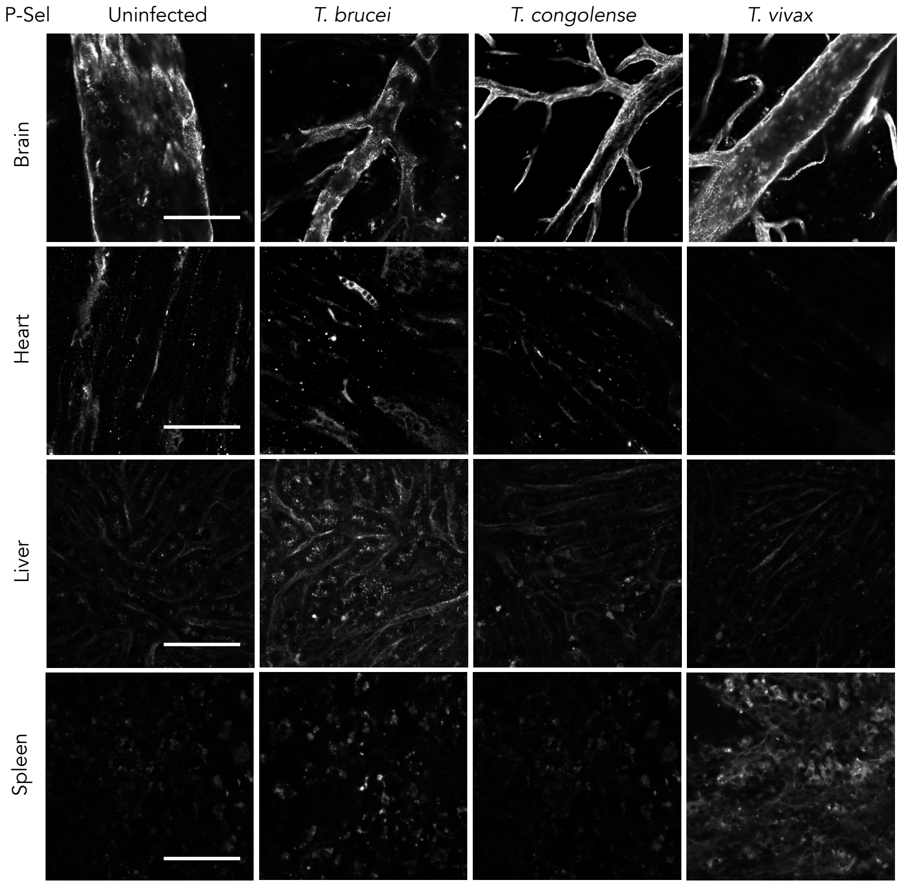


**Supplementary Figure 7. Expression P-selectin at the first peak of parasitaemia of African trypanosome infections in C57B/6J mice.** Representative images of P-selectin in the brain, heart, liver and spleen, detected by fluorescence microscopy. Scale bar is 40 μm.

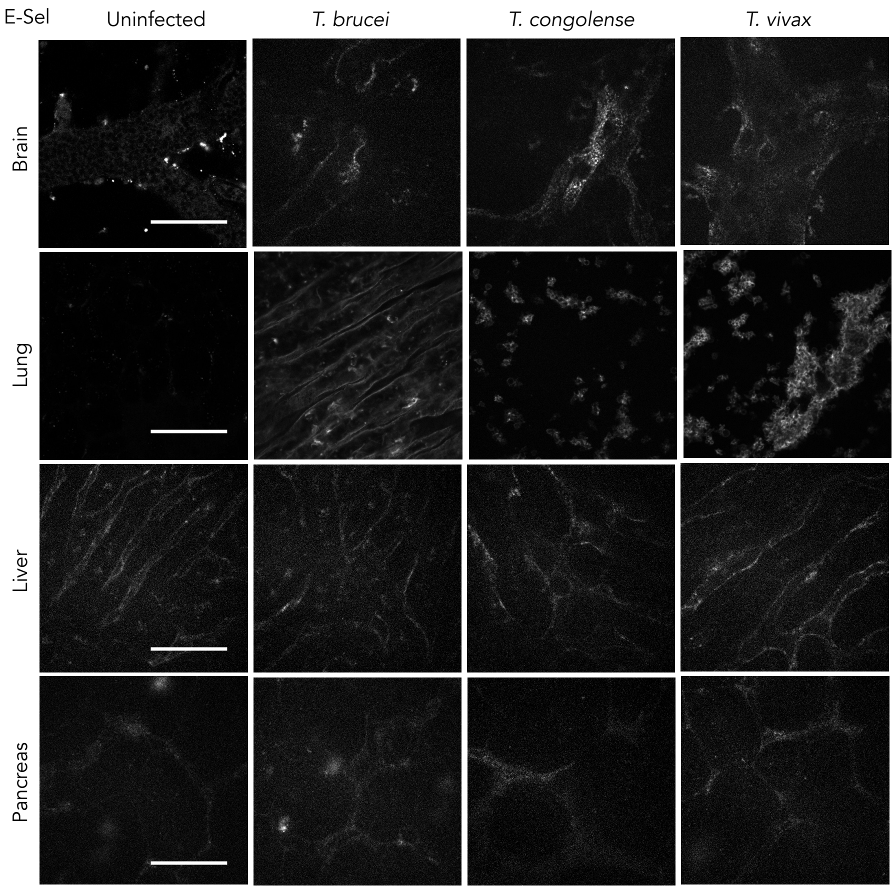


**Supplementary Figure 8. Expression E-selectin at the first peak of parasitaemia of African trypanosome infections in C57B/6J mice.** Representative images of E-selectin in the brain, lungs, liver and pancreas, detected by fluorescence microscopy. Scale bar is 40 μm.

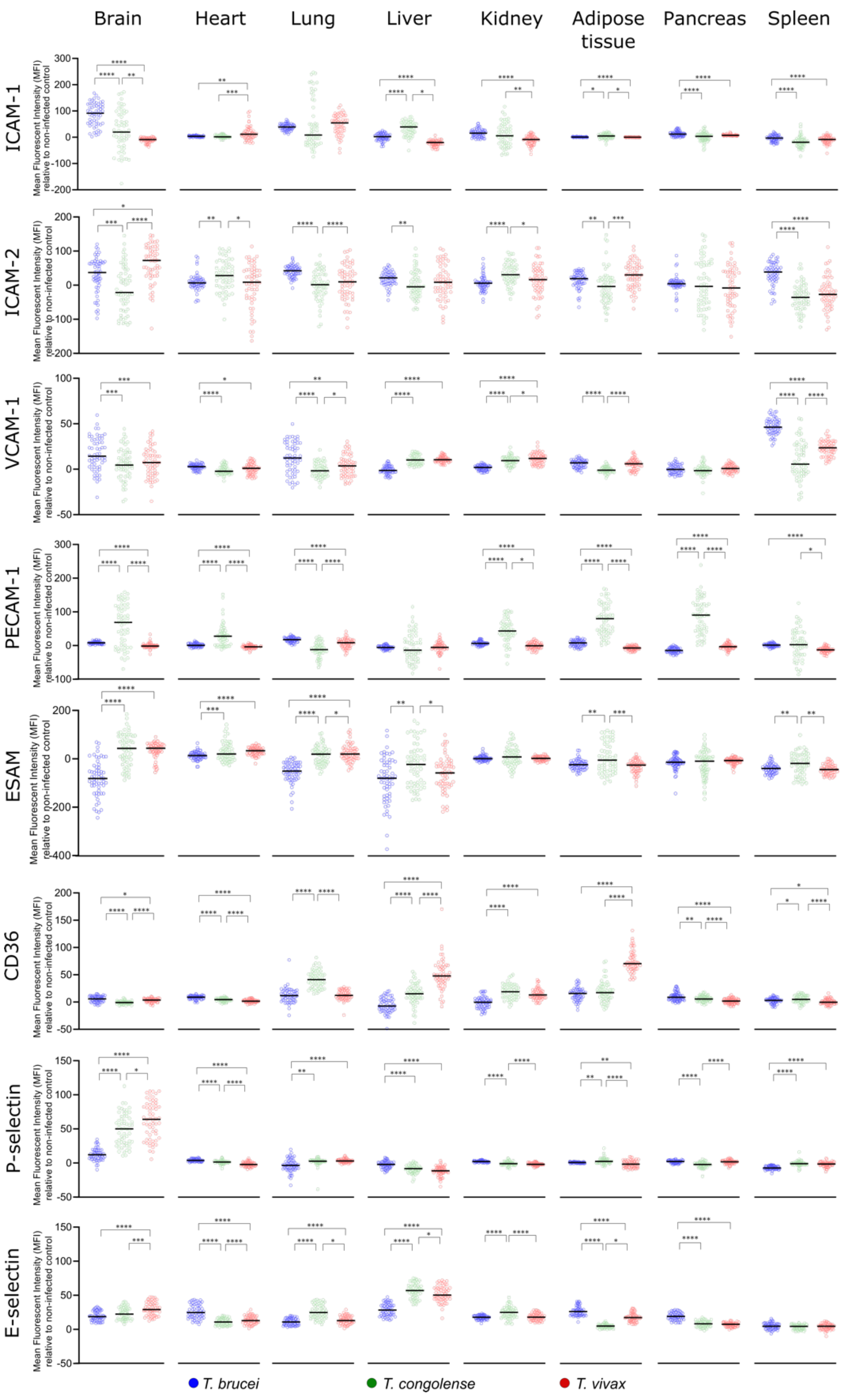


**Supplementary Figure 9. Quantification of expression changes of each endothelial cell surface protein in the vasculature of major organs, detected by fluorescence microscopy.** Mean fluorescence intensity values of each endothelial cell receptor for individual vessels were compared between species. Repeated Measures, One-way ANOVA, with Tukey's Multiple Comparisons test. * p-value < 0.05; ** p-value < 0.01; *** p-value < 0.001; **** p-value < 0.0001.

### Supplementary Video Legends

**Supplementary Video 1.** *T. vivax* in the brain vasculature of an infected C57B/6 mouse. Mice were intravenously injected with Hoechst 33342 and 70 kDa FITC-Dextran. Hoechst enables visualization of *T. vivax* nuclei and kinetoplasts. Contrast generated by FITC-Dextran enables visualization of the parasite body. Images were obtained at a rate of 5 frames per second.

**Supplementary Video 2.** *T. vivax* in the pancreas vasculature of an infected C57B/6 mouse. Mice were intravenously injected with Hoechst 33342 and 70 kDa FITC-Dextran. Contrast generated by FITC-Dextran enables visualization of the parasite body, including flagellar beating and its interaction with the host vasculature. Images were obtained at a rate of 5 frames per second.

**Supplementary Video 3.** *T. vivax* in the pancreas vasculature of an infected C57B/6 mouse. Mice were intravenously injected with Hoechst 33342 and 70 kDa FITC-Dextran. Contrast generated by FITC-Dextran enables visualization of the parasite body, including flagellar beating and its interaction with the host vasculature. Blood flow was blocked by capillary ligation to enable observation of parasite displacement (or lack thereof). Images were obtained at a rate of 5 frames per second.

**Supplementary Video 4.** *T. vivax* in the pancreas vasculature of an infected C57B/6 mouse. Mice were intravenously injected with Hoechst 33342 and 70 kDa FITC-Dextran. Contrast generated by FITC-Dextran enables visualization of the parasite body, including flagellar beating and its interaction with the host vasculature. Blood flow was blocked by capillary ligation to enable observation of parasite displacement (or lack thereof). Images were obtained at a rate of 5 frames per second.

**Supplementary Video 5.** Close up video of *T. vivax* in the heart vasculature of an infected C57B/6 mouse. Mice were intravenously injected with Hoechst 33342 and 70 kDa FITC-Dextran. Contrast generated by FITC-Dextran enables visualization of the parasite body, including flagellar beating and its lack of displacement under flow. Images were obtained at a rate of 5 frames per second.
